# Quantitative live cell imaging of a tauopathy model enables the identification of a polypharmacological drug candidate that restores physiological microtubule regulation

**DOI:** 10.1101/2022.10.31.514565

**Authors:** Luca Pinzi, Christian Conze, Nicolo Bisi, Gabriele Dalla Torre, Nanci Monteiro-Abreu, Nataliya I. Trushina, Ahmed Soliman, Andrea Krusenbaum, Maryam Khodaei Dolouei, Andrea Hellwig, Michael S. Christodoulou, Daniele Passarella, Lidia Bakota, Giulio Rastelli, Roland Brandt

## Abstract

Tauopathies such as Alzheimer’s disease are characterized by the aggregation and increased phosphorylation of the microtubule-associated protein tau. The pathological changes in tau are closely linked to neurodegeneration, making tau a prime candidate for intervention. However, the multiple facets of tau function and the lack of cellular tauopathy models that could support mechanism-based drug development hampers progress. Here we report the development of a live-cell imaging approach to quantitatively monitor pathological changes of human tau as it interacts with axonal microtubules. We show that a full-length aggregation-prone tau construct exhibits reduced interaction with microtubules as it increasingly aggregates. Through chemoinformatic analyses, we identified 2-phenyloxazole (PHOX) derivatives as putative polypharmacological small molecules that inhibit tau aggregation and modulate tau phosphorylation. We found that PHOX15 restores the physiological microtubule interaction of aggregation-prone tau in neurons and inhibits the first phase of tau aggregation *in vitro*. Furthermore, we report that PHOX15 inhibits the tau kinases GSK3β and Cdk5, alters the kinome activity of model neurons, and reduces tau phosphorylation at disease-relevant sites. Molecular dynamics simulations highlight cryptic channel-like pockets crossing tau protofilaments and indicate that the binding of PHOX15 in one of the channels reduces the protofilament’s ability to adopt a PHF-like conformation. The data show that our imaging approach provides a useful tool for identifying compounds that modulate tau-microtubule interaction in axons. We demonstrate that a polypharmacological approach to simultaneously treat tau aggregation and tau phosphorylation is able to restore physiological microtubule regulation, identifying PHOX15 as a promising drug candidate to counteract tau-induced neurodegeneration.

## INTRODUCTION

Tauopathies are a group of neurodegenerative disorders associated with the accumulation of abnormal tau protein in the brain (Arendt et al., 2016). The most common tauopathy is Alzheimer’s disease (AD), in which intracellular aggregates of tau with increased phosphorylation (“hyperphosphorylation”) are joined by extracellular amyloid plaques containing aggregated amyloid-*β* (A*β*) (Selkoe and Hardy, 2016). Because tau mutations are sufficient to cause tauopathies and because tau inclusions correlate much better with cognitive impairment than amyloid plaques do, tau pathology is considered to be the major driver for neuronal degeneration in AD and other tauopathies (Clark et al., 1998; Hutton et al., 1998; Nelson et al., 2012; Spillantini et al., 1998).

Due to the failure of A*β*-targeted therapies, tau has become a target of rapidly evolving therapeutic strategies (Chang et al., 2021; Soliman et al., 2022). However, tau is a challenging target due to its complex interactions as an intrinsically disordered protein, its various post-translational modifications, and the complexity of tau pathologies (Brandt et al., 2020; Limorenko and Lashuel, 2021; Shi et al., 2021). Mechanisms of toxicity are also controversial, but prefibrillar tau oligomers and soluble tau with disease-like modifications can be toxic species (Fath et al., 2002; Patterson et al., 2011). Furthermore, disruption of the physiological function of tau, particularly its dynamic interaction with axonal microtubules, can be a crucial contributor to the disease (Conze et al., 2022).

Because abnormal forms of tau could trigger a plethora of pathomechanisms, targeting individual downstream mechanisms may have limited therapeutic effects. Polypharmacological drugs may provide a solution to this problem by simultaneously modulating tau aggregation, tau phosphorylation, and potential additional tau functions that may be impaired due to disease. However, cell-based models of tauopathies that would help to monitor tau activity and behavior in neurons and to develop mechanism-based therapies are scarce. Previously, biosensor cells to monitor tau aggregation have been generated (Kfoury et al., 2012), however, the cell lines are based on the expression of artificial tau fragments, and the relevance for the formation of authentic amyloidogenic tau aggregates has been questioned (Kaniyappan et al., 2020). In addition, since the tau regions involved in aggregation and microtubule-binding overlap, it would be important to determine whether tau aggregation inhibitors that bind covalently or non-covalently to the microtubule-binding region of tau (Cisek et al., 2014) interfere with the physiological interaction of tau with microtubules.

In this work, we developed a live-cell imaging assay to identify compounds that restore physiological microtubule interaction of an aggregation-prone human tau construct in axon-like processes of model neurons and axons of primary neurons. Using a panel of small molecules predicted to have tau and kinase modulating activity, we identified the 2-phenyloxazole derivative PHOX15, which restores the physiological tau-microtubule interaction in cells. We show that this small molecule blocks the first phase of tau aggregation *in vitro*, inhibits the tau kinases GSK3*β* and Cdk5, and reduces tau phosphorylation at selected pathogenic sites in the proline-rich region of tau. Molecular dynamics (MD) simulations identified cryptic channel-like pockets crossing tau protofilaments not previously described. The simulations suggest that the binding of PHOX15 in one of these channels reduces the ability of the protofilament to adopt a PHF-like conformation, consistent with PHOX15 exerting an inhibitory effect in the early stages of tau oligomerization.

## RESULTS

### Aggregation-prone human tau exhibits reduced microtubule interaction in neuronal cells and progressively aggregates in neurons

We used the single amino acid deletion mutant TauΔK280 found in a case of frontotemporal dementia (FTD) and AD (Momeni et al., 2009; Rizzu et al., 1999) to develop a cellular assay for compounds that affect the tau-microtubule interaction. *In vitro* experiments had previously demonstrated that this mutant showed an increased propensity to form paired helical filaments (Barghorn et al., 2000). Consistent with this, recombinant human TauΔK280 showed greater than 50% increased aggregation compared to wild-type tau in heparin-induced cell-free aggregation assays (**Figure 1A**). In standard *in vitro* microtubule-polymerization assays, wild-type tau and TauΔK280 showed very similar activity in promoting microtubule assembly, consistent with little or no tau aggregation under cell-free conditions in the absence of heparin (data not shown).

**Figure 1:**
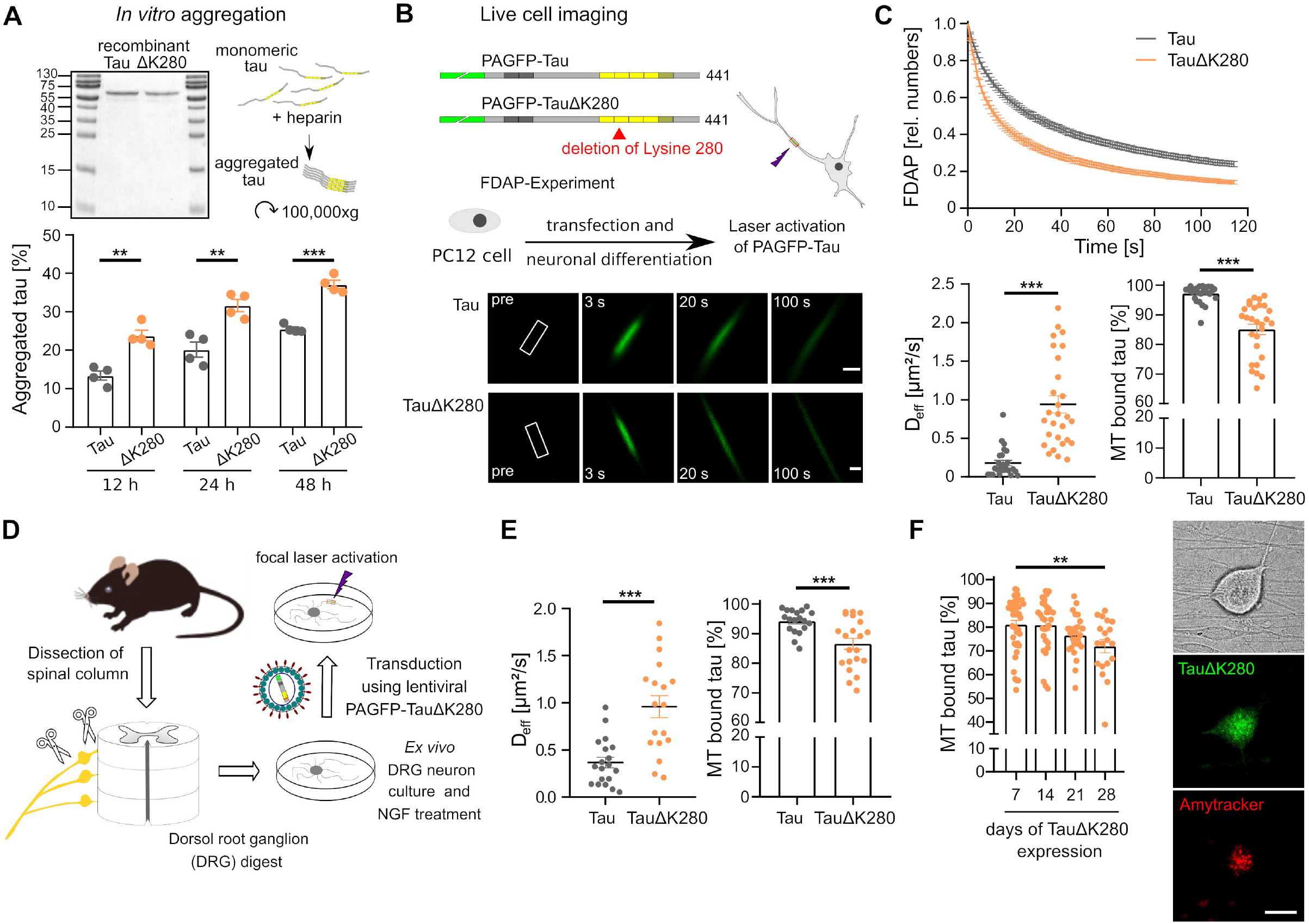
Aggregation-prone human tau exhibits reduced microtubule interaction in neuronal cells and progressively aggregates in neurons. **A**. *In vitro* aggregation of recombinant human wild-type tau (Tau) and aggregation-prone tau (TauΔK280). Coomassie Brilliant Blue-stained SDS-PAGE of the purified proteins (top left) and fractions of aggregated tau determined by ultracentrifugation after incubation with heparin for the indicated times are shown (bottom; mean ± SEM; n = 4). Molecular mass standards are indicated. Statistically significant differences between Tau and TauΔK280 as determined by an unpaired Student’s t-test are displayed. ** p <0.01; *** p <0.001. **B**. Live cell imaging of tau dynamics in axon-like processes of neuronally differentiated PC12 cells. Schematic representations of the expressed tau constructs are shown above. The MT-binding repeat regions (RR1–RR4) are indicated by yellow boxes, the pseudorepeat region in dark yellow, and the N-terminal PAGFP fusion in green. Adult-specific exons in the N-terminus of tau (N1, N2) are shown in dark gray. The lysine deletion of the aggregation-prone construct is in the second repeat. Shown below are representative time-lapse micrographs of a fluorescence decay after photoactivation (FDAP) experiment. Scale bar, 3 µm. **C**. FDAP diagrams after photoactivation of PAGFP-Tau or PAGFP-TauΔK280-expressing PC12 cells. Mean ± SEM of 26 (Tau) and 28 (TauΔK280) cells are shown. Scatterplots of effective diffusion constants (D_eff_) and percentage of tau bound to microtubules are shown below. Statistically significant differences between Tau and TauΔK280 as determined by unpaired Student’s t-tests with Welch correction are indicated. *** p <0.001. **D**. Preparation of primary neurons from dorsal root ganglia (DRG). DRG neurons were prepared from adult mice by extracting the ganglia delineating the spinal cord and sequential enzymatic digestion. Neurons were lentivirally transduced to continuously express PAGFP-Tau or PAGFP-TauΔK280. **E**. Scatterplots of effective diffusion constants (D_eff_) (left) and percentage of tau bound to microtubules (right) in axons of DRG neurons. Mean ± SEM of 19 (Tau) and 22 (TauΔK280) infected cells are shown. Statistically significant difference between Tau and TauΔK280 as determined by unpaired Student’s t-tests with Welch correction are indicated. *** p <0.001. **F**. Effect of continuous TauΔK280 expression on the percentage of tau bound to microtubules in axons of DRG neurons. Micrographs of a DRG neuron after 3 weeks of TauΔK280 expression showing a transmitted light micrograph, PAGFP-TauΔK280 fluorescence, and the Amytracker 680 signal of a live DRG neuron are displayed on the right. Mean ± SEM are shown. Statistically significant differences from the 7-day value determined by ANOVA with Dunnet’s post hoc test are indicated. ** p <0.01. Scale bar, 20 µm.

We used fluorescence decay after photoactivation (FDAP) experiments to determine a possible change in the interaction of TauΔK280 with microtubules in axon-like processes of model neurons. The constructs were N-terminally tagged with photoactivatable GFP (PAGFP) and exogenously expressed in PC12 cells, which were differentiated into a neuronal phenotype. After focal activation, both constructs showed dissipation from the region of activation, which appeared higher for TauΔK280 (**Figure 1B**). Indeed, the FDAP curves showed increased decay of TauΔK280 compared to wild-type tau, indicating reduced interaction with microtubules in the cellular environment (**Figure 1C, top**). Accordingly, calculation of the effective diffusion constant using a one-dimensional diffusion model resulted in a significantly higher value for TauΔK280. This corresponds to a >10% reduced binding to microtubules of TauΔK280 (**Figure 1C, bottom**), indicating a reduced availability of the aggregation-prone tau to interact with microtubules.

To determine whether TauΔK280 exhibits reduced microtubule interaction also in authentic axons of neurons, we transduced primary dorsal root ganglia (DRG) neurons dissected from adult mice to express the respective PAGFP-tagged tau constructs (**Figure 1D**). Again, TauΔK280 showed higher effective diffusion and a very similar decrease in axonal microtubule-binding of TauΔK280 compared to wild-type-tau, consistent with a reduced availability of the aggregation-prone tau construct also in primary neurons (**Figure 1E**). The impaired microtubule interaction of TauΔK280 is likely caused by the formation of soluble tau oligomers in the cells. Of note, the tau oligomers were small enough not to cause reduced diffusion in our assay system. To test the hypothesis that the soluble tau oligomers are precursors to the formation of larger, insoluble tau aggregates, we analyzed DRG neurons several weeks after lentiviral infection to prolong the presence of the aggregation-prone TauΔK280 in the cells. Indeed, three weeks post-infection, tau aggregates could be observed in most cell bodies. The tau aggregates were positive after live staining with Amytracker (Ebba Biotech, Sweden), indicating formation of tau amyloid with more than eight parallel *β*-sheets in register. Parallel to aggregate formation, binding of TauΔK280 to microtubules progressively decreased with expression time (**Figure 1F**).

The results demonstrate a reduced microtubule interaction of TauΔK280 in axon-like processes of model neurons and axons of primary neurons. The time course of TauΔK280-expression in primary neurons indicates that the reduced microtubule interaction is caused by the formation of small soluble tau oligomers as precursors to tau amyloids that form after prolonged tau exposure. The data also point out that this imaging approach provides a useful tool to identify compounds that modulate tau oligomerization and tau-microtubule interaction in axons of living neurons.

### Chemoinformatic analyses indicate that PHOX compounds can inhibit tau aggregation

We have previously identified 2-phenyloxazole (PHOX) derivatives as selective monoamine oxidase B (MAO-B) inhibitors (Di Paolo et al., 2019) (**Figure 2A**). MAO-B inhibitors are used to treat Parkinson’s disease and are also considered potential drug candidates for the treatment of AD (Jost, 2022; Park et al., 2019). In accordance with the concept of polypharmacology, i.e., the use of a single drug acting on multiple targets or disease pathways (Anighoro et al., 2014), we searched for possible additional targets of the PHOX compounds through integrated 2D (MACCS and ECFP4 fingerprints; OpenEye Toolkits 2020.2.2 OpenEye Scientific Software, Santa Fe, NM. Available online: http://www.eyesopen.com) and 3D (ROCS, OpenEye) (Hawkins et al., 2007) computational similarity estimations.

**Figure 2:**
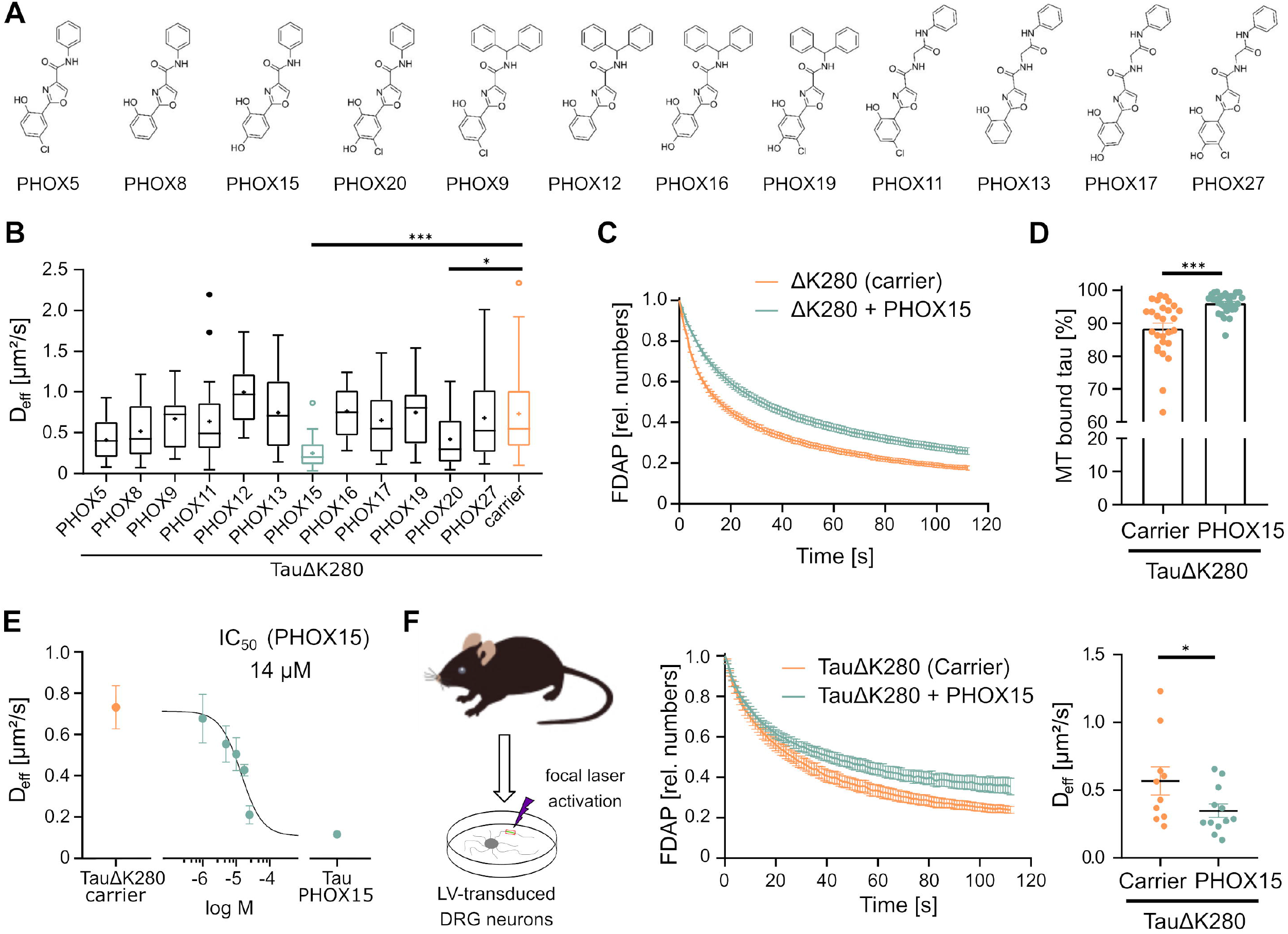
PHOX15 restores the physiological microtubule interaction of aggregation-prone tau. **A**. Chemical structures of the investigated PHOX compounds. **B**. Boxplots of effective diffusion constants (D_eff_) in response to the twelve PHOX derivatives shown in Figure 2. TauΔK280-expressing neuronally differentiated PC12 cells were treated with either 25 µM compound or carrier (0.125% DMSO) for 20 hours prior to imaging. Tukey plot with median (-) and mean (+) of 15-30 cells per condition is shown. Statistically significant differences in compound to carrier were determined by one-way ANOVA with Dunnett’s post hoc. * p <0.05; *** p <0.001. **C**. FDAP plots after photoactivation of PAGFP-TauDK280-expressing PC12 cells with the most active compound (PHOX15) or carrier (0.125% DMSO). Mean ± SEM of 30 (PHOX15) and 27 (carrier) cells are shown. **D**. Scatterplot of percentage of tau bound to microtubules based on the data shown in (B). Statistically significant difference determined by unpaired Student’s t-tests with Welch correction is indicated. *** p <0.001. **E**. Dose-response curve of the effect of PHOX15 on the effective diffusion constants (D_eff_) of TauΔK280-expressing PC12 cells. Means ± SEM of 15-20 cells per concentration (1-25 µM) are shown. The curve was fitted using a four-parameter log-logistic model. The plateau of the curve was constrained to D_eff_ of TauΔK280 with carrier (0.125% DMSO, n=27) and Tau with 25 µM PHOX15 (n=26) according to (Weimer et al., 2012). **F**. FDAP plots and scatterplot of effective diffusion constants (D_eff_) after photoactivation of lentivirally transduced DRG neurons expressing PAGFP-TauΔK280. The effect of PHOX15 (25 µM, treatment of 20 hours prior to imaging) compared to a carrier control (0.125% DMSO) is shown (Meana ± SEM from 12 (PHOX15) and 10 cells (control) are shown. Statistically significant differences between samples determined by an unpaired Student’s t-test are indicated. * p <0.05.

The 2D and 3D similarity analyses allowed us to prioritize a set of potential targets of the PHOX compounds according to their similarity to active ligands reported in ChEMBL (Gaulton et al., 2017). Remarkably, the microtubule-associated protein tau was one of the best-scoring targets emerging from the consensus of the 2D and 3D methods. In particular, PHOX 5, 8, 15 and 20 were closely related to potent tau aggregation inhibitors. Representative 3D alignments show high conformational overlap and significant superposition of hydrogen-bond acceptor/donor and aromatic features.

We further investigated whether PHOX compounds might possess molecular properties and scaffolds required for tau anti-aggregation activity (Pinzi et al., 2021). The vast majority of the PHOX compounds met these requirements, with the exception of the bulkier benzylbenzene derivatives PHOX 9, 12, 16, and 19. In particular, the molecular properties of PHOX 5, 8, 11, 13, and 15 were similar to those of highly active tau ligands. These compounds have an amide group, three aromatic rings (one of which being a heterocycle), around six hydrogen-bond acceptors and a low aliphatic character (Pinzi et al., 2021). Finally, the compounds were tested *in silico* for their predicted ability to cross the Blood-Brain Barrier (BBB) by using the *QikProp* software (Schrödinger Release 2022-1: QikProp; Schrödinger, LLC: New York, NY, USA, 2022).

The results of the chemoinformatic analyses suggest that the PHOX compounds can affect tau oligomerization and possibly cross the BBB, two important features that make these compounds worthy of further investigation.

### The 2-phenyloxazole derivative 2-(2,4-dihydroxyphenyl)-N-phenyloxazole-4-carboxamide (PHOX15) restores the physiological microtubule interaction of aggregation-prone tau in neurons

The twelve PHOX derivatives shown in **Figure 2A** were tested for their effect on metabolic activity and cytotoxicity profiles on differentiated model neurons. While some PHOX derivatives affected MTT conversion in a concentration-dependent manner, indicating biological activity, none of the compounds were toxic at concentrations of 25 µM or below.

Next, we used our live cell imaging approach to test the PHOX derivatives for their activity to modulate tau-microtubule interaction at subtoxic concentrations. Two of the compounds (PHOX15 and PHOX20) significantly reduced the effective diffusion constant, with PHOX15 being the most efficient (**Figure 2B**). Indeed, the FDAP curves showed a significantly reduced dissipation of TauΔK280 in the presence of PHOX15 compared to a vehicle control (**Figure 2C**), indicating increased microtubule interaction. Accordingly, the calculation of the amount of microtubule-bound tau showed an ∼10% increased binding to microtubules, almost to the level of wild-type tau protein (**Figure 2D**). We also determined the IC_50_ value of the activity of PHOX15 to restore microtubule binding of TauΔK280, obtaining a value of 14 µM, which is in the concentration range of the expressed tau construct (**Figure 2E**). The results suggest that PHOX15 binds to pathological tau, inhibits tau oligomerization, and restores the physiological interaction between tau and microtubules in axon-like processes of neuronal cells.

In addition, we examined the activity of PHOX15 in axons of DRG neurons infected to express TauΔK280. Also, in authentic axons, treatment with PHOX15 reduced the effective diffusion constant to that of neurons infected to express wild-type tau, indicating that PHOX15 is able to restore the physiological interaction of aggregation-prone tau with microtubules in primary neurons (**Figure 2F**).

### PHOX15 inhibits the first phase of tau aggregation

To determine the mechanism by which PHOX15 may interfere with tau aggregation, we performed heparin-induced cell-free aggregation assays in the presence or absence of PHOX15. Recombinant TauΔK280 assembled into single filament structures with a common helical turn period very similar to what was previously observed for cylindrical filaments derived from paired helical filaments (PHFs) (Ruben et al., 2005). The filaments had a diameter of 9-12 nm and a helical turn period (distance between the thin regions) of 70-80 nm (**Figure 3A**). The presence of PHOX15 did not affect the structure of the filaments to any major extent, and the diameter of the cylindrical filaments and the turn period were similar.

**Figure 3:**
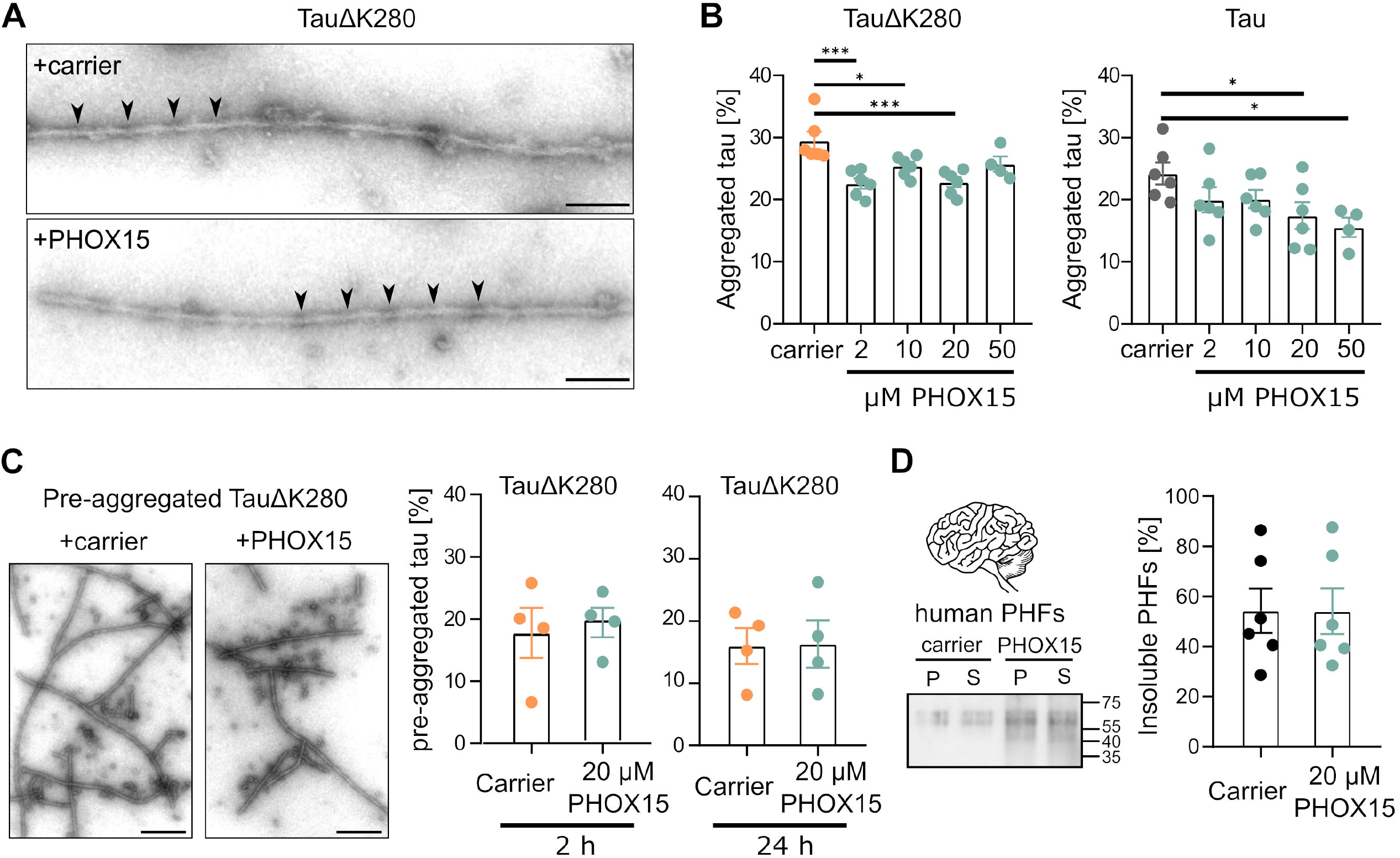
PHOX15 inhibits the first phase of tau aggregation. **A**. Electron micrographs of tau filaments that were formed in the presence of heparin from purified TauΔK280 (0.4 mg/ml) in the presence of PHOX15 (20 µM) or carrier (1% DMSO). Samples were incubated for 24 hrs at 37°C, and negatively stained with 1% uranyl acetate. Scale bar, 100 nm. Thin regions of the cylindrical filaments marked by arrowheads indicate a helical turn period of 70-80 nm. **B**. Effect of PHOX15 on *in vitro* aggregation of recombinant aggregation-prone tau (TauΔK280) and human wild-type tau (Tau). Fractions of aggregated tau determined by ultracentrifugation after incubation for 24 hrs with the indicated concentration of PHOX15 or carrier (1% DMSO) are shown (mean ± SEM; n = 4-6). Statistically significant differences between PHOX15-treated samples and control as determined by an unpaired Student’s t-test are displayed. * p <0.05; *** p <0.001. **C**. Effect of PHOX15 on pre-aggregated filaments prepared from TauΔK280. Electron micrographs of tau filaments (24 hrs pre-aggregated) treated with PHOX15 (20 µM) or carrier (1% DMSO) for additional 24 hrs. Fractions of aggregated tau determined by ultracentrifugation are shown on the right. **D**. Effect of PHOX15 on PHFs from human brain. Samples of 0.1 mg/ml SDS-soluble PHFs (Ksiezak-Reding et al., 1994) were treated for 24 hrs with PHOX15 (20 µM) or carrier (1% DMSO). Immunoblot of the pellet (P) and (S) supernatant fractions of the PHFs stained with anti-PHF1 antibody (Greenberg et al., 1992) (left panel). Molecular mass standards are indicated. Fractions of insoluble PHFs determined by ultracentrifugation are shown on the right (mean ± SEM; n = 4).

However, PHOX15 reduced filament formation by approximately 20%, as quantified by sedimentation assays (**Figure 3B**). When added to preformed tau aggregates, PHOX15 had no effect on the structure and amounts of filaments (**Figure 3C**). Consequently, treatment with PHOX15 also did not decrease the amount of SDS-soluble PHFs isolated from human brain (Ksiezak-Reding et al., 1994) (**Figure 3D**).

Cylindrical filaments with periodically thin regions are considered to be the PHF precursor filament (Ruben et al., 2005). Thus, the results indicate that PHOX15 inhibits the first phase of tau aggregation, likely by blocking tau oligomerization or protofibril formation.

### PHOX15 inhibits the tau kinases GSK3β and Cdk5, changes the kinome activity of the cells, and reduces tau phosphorylation at disease-relevant sites

Aberrant phosphorylation and aggregation are key features of tau in the brains of AD patients, and high stoichiometric increased tau phosphorylation (hyperphosphorylation) is thought to result in tau dysfunction and pathological properties (Arendt et al., 2016; Trushina et al., 2019). Several kinases have been implicated in the aberrant phosphorylation of tau and are thought to contribute to disease progression. Of these, glycogen synthase kinase 3β (GSK3β) and cyclin-dependent kinase 5 (Cdk5) have been implicated in AD (Kimura et al., 2014; Lauretti et al., 2020).

Notably, GSK3β also emerged as a top-ranked target in the similarity estimates, adding further strength to the putative polypharmacological behavior of this ligand. Docking of PHOX15 based on available crystal structures suggests that the binding mode involves an interaction with the ATP pocket of GSK3β (**Figure 4A**). In the complex, the amide carbonyl hydrogen bonds with the backbone nitrogen of the hinge, the phenyl ring occupies the hydrophobic pocket of the kinase, and the para-hydroxyl group hydrogen bonds with both the Glu56 residue of the *α*C helix and the Asp157 residue of the highly conserved Asp-Phe-Gly (DFG) motif. This binding mode is consistent with the structure-activity relationships (SAR) of other PHOX derivatives tested in this work (data not shown). *In vitro* kinase assays on these targets showed that PHOX15 did inhibit both GSK3β and Cdk5 activity with IC_50_ values of 1.9 and 1 µM, respectively (**Figure 4B**), confirming the predictions.

**Figure 4:**
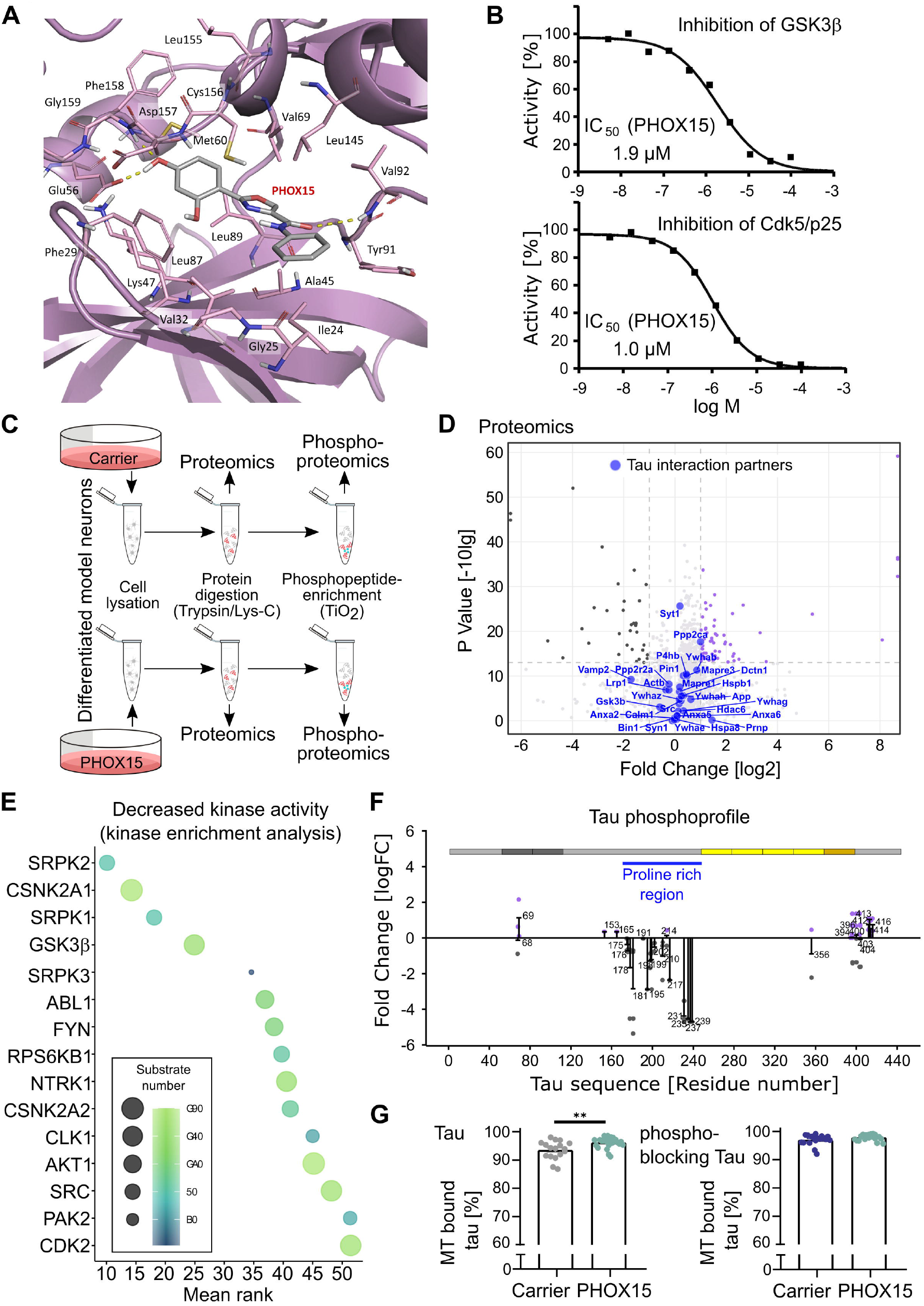
PHOX15 inhibits the tau kinases GSK3β and Cdk5, changes the kinome activity of the cells, and reduces tau phosphorylation at disease-relevant sites. **A**. Binding mode of PHOX15 at the ATP pocket of GSK3β based on docking analysis using available crystal structures. **B**. *In vitro* kinase assays for inhibition of GSK3β and Cdk5 activity by different concentrations of PHOX15. **C**. Schematic representation of the approach for proteomics and phosphoproteomics analyses of differentiated model neurons treated for 20 hrs with PHOX15 (25 µM) or carrier (0.125% DMSO). **D**. Volcano plot showing up- or down-regulated proteins in PHOX15-treated cultures compared to controls. None of the >60 experimentally confirmed tau interaction partners (Trushina et al., 2019) were significantly up-or down-regulated. **E**. Kinase enrichment analysis of the phosphoproteomic data to identify the pattern of kinases responsible for increased phosphorylation of the cellular proteins as response to PHOX15 treatment. **F**. Phosphoprofile of endogenous tau as determined by phosphoproteomics. The changes in the phosphorylation of individual sites as a response to treatment with PHOX15 are displayed. **G**. Scatterplots showing percentage of wild-type tau and phosphoblocking tau bound to microtubules as determined by FDAP experiments with the respective PAGFP-tagged tau constructs. Cells were treated with PHOX15 (25 µM) or carrier (0.125% DMSO) for 20 hrs prior to imaging. Mean ± SEM of 26, 17 and 22, 18 (carrier) for wild-type tau and phosphoblocking tau expressing cells, respectively, are shown. Statistically significant differences as determined by unpaired Student’s t-tests with Welch correction are indicated. ** p <0.01.

To systematically determine the effect of PHOX15 on gene expression and protein phosphorylation, we performed proteomics and phosphoproteomics analyses of model neurons treated with PHOX15 or a vehicle (**Figure 4C**). None of the >60 experimentally confirmed tau interaction partners (Trushina et al., 2019) was significantly down-or up-regulated in PHOX15-treated neurons (**Figure 4D**). This suggests that the changes in tau aggregation and tau-microtubule interaction are independent of the differential expression of components of the tau interactome.

To determine the effect of PHOX15 on the kinome activity of the cell, we performed a kinase enrichment analysis to link the identified phosphorylation sites to the reduced activity of the kinases that are most likely responsible for the decreased protein phosphorylation (Kuleshov et al., 2021; Lachmann and Ma’ayan, 2009). GSK3β ranked fourth in the list (**Figure 4E**), indicating that PHOX15 affected GSK3β-dependent phosphorylation in neural cells as well.

Next, we determined the effect of PHOX15 on the phosphoprofile of endogenous tau by phosphoproteomics. Phosphorylation of PHOX15-treated neurons was greatly reduced at several serine and threonine residues in the proline-rich region (PRR) of tau aminoterminally flanking the microtubule-binding region (**Figure 4F**). Tau’s PRR is considered to be a signalling module that regulates tau’s intracellular interactions, including its activity to interact with microtubules (Brandt et al., 2020). The PRR has the highest relative content of serine/threonine residues, making it the major region of tau for phosphorylation (Brandt et al., 2020). Phosphorylation of multiple sites, particularly a phosphocluster between Thr231 and Ser239, showed a large decrease in phosphorylation after PHOX15 treatment. This phosphocluster includes major phosphorylation sites identified in PHFs from patients with AD (Thr231, Ser235) (Morishima-Kawashima et al., 1995), which are predicted phosphorylation sites for GSK3β and Cdk5. Other sites of severe reduction include Thr181, an established core biomarker for cerebrospinal fluid (CSF) in AD (Vanmechelen et al., 2000), which is also predicted to be phosphorylated by GSK3β and Cdk5 (NetPhos-3.1b prediction (Blom et al., 2004)). Notably, Thr231 was recently identified as a master site governing the propagation of tau phosphorylation at several AD-associated epitopes (Stefanoska et al., 2022).

To determine whether PHOX15 increases microtubule interaction by reducing tau phosphorylation at disease-relevant sites, we performed FDAP assays to assess a possible change in the interaction of wild-type tau and a phosphoblocking tau construct with microtubules in axon-like processes of model neurons. The phosphoblocking tau construct was designed by mutating to alanine the ten major phosphorylation sites previously identified as hyperphosphorylated in tau from AD patients (Morishima-Kawashima et al., 1995; Trushina et al., 2019), to prevent phosphorylation at these residues. Five of these sites were in the PRR (Ser198, Ser199, Ser202, Thr231, Ser235); all showed reduced phosphorylation as a result of PHOX15 treatment (**Figure 4F**). Treatment with PHOX15 increased microtubule association of wild-type tau but failed to do so with phosphoblocking tau (**Figure 4G**). This finding suggests that PHOX15 has a dual effect on tau-microtubule interaction. First, it restores the physiological interaction of an aggregation-prone construct by reducing tau aggregation, and second, it increases microtubule interaction by reducing tau phosphorylation at disease-relevant sites.

### Molecular dynamics simulations highlight cryptic channel-like pockets crossing tau protofilaments

The cryo-EM structure of the tau filaments from AD brains (PDB ID: 5O3L) reveals a C-shaped structure composed of stacked tau sequences encompassing residues 306-378 of tau’s microtubule-binding domain (Falcon et al., 2018; Fitzpatrick et al., 2017) (**Figure 5A, left**). The two protofilaments A and B form a dimeric structure that can assemble into a PHF conformation (**Figure 5A, right**). Given that the effect of PHOX15 was observed at an early stage of tau aggregation, we investigated how the molecule might influence and impair protofilament formation. An 820 ns classical MD simulation of a single tau protofilament in water showed that the structure of the protofilament remained relatively stable during the MD simulation, with minor variations in Root Mean Squares Deviation (RMSD), the radius of gyration, β-sheet content and intra-molecular H-bond pattern. The analyses revealed the presence of three cryptic pockets with channel-like features (**Figure 5B**; pocket 1 (P1) formed by residues Asp314, Glu372, His374, Lys370, Pro312 and Val313; P2 by residues Asn359, Asp358, Gly333-335, Leu357, Pro332, and P3 by residues Ser320, Lys321, Cys322, Gly323, Val363, Pro364, Gly365, and Gly366). The pockets were not present in the initial cryo-EM structure 5O3L (Fitzpatrick et al., 2017) and to the best of our knowledge have not been described previously.

**Figure 5:**
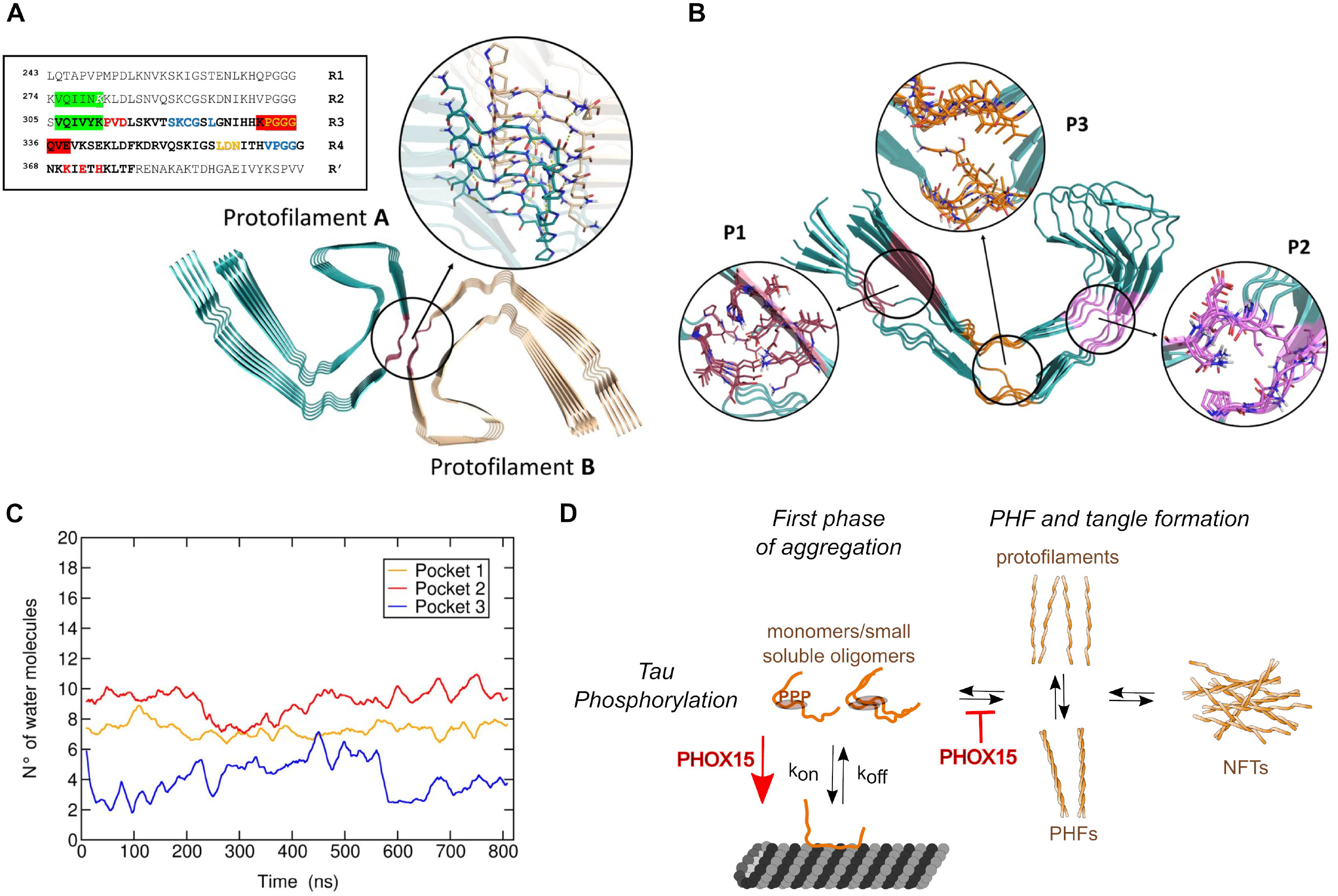
Molecular dynamics simulations highlight cryptic channel-like pockets crossing tau protofilaments. **A**. Cryo-EM structure of the PHF dimer. Glycine triads are highlighted in red. A close-up view of the glycines interacting between the protofilaments of the PHF is also shown. The sequence of the MT-binding repeat region of the longest Tau isoform containing four repeats (R1, R2, R3 and R4) is shown on the right. Residues elucidated by cryo-EM (PDB ID: 5O3L) (Fitzpatrick et al., 2017), which form the PHF core are shown in bold. Aggregation-prone sequences PHF6 and PHF6* are highlighted in blue and the key lysine residue of the aggregation-prone ΔK280 mutant is shown in white italic character. Residues that form the protofilament-protofilament dimerization site are highlighted in red. Residues forming pocket 1, pocket 2 and pocket 3 are shown in red, yellow and blue character, respectively. **B**. Snapshot of protofilament A after 200 ns of MD simulation. Pocket 1, pocket 2 and pocket 3 are highlighted. **C**. Time evolution of the number of water molecules permeating pocket 1, pocket 2 and pocket 3. Standard deviations are not shown for clarity. **D**. Model showing the effect of PHOX15 on tau aggregation. Our data indicate that PHOX15 inhibits the first phase of tau aggregation and decreases the phosphorylation of tau. Both activities increase the interaction of tau with microtubules to a physiological level.

The pockets present channel-like features with a non-regular shape, as their cavities cross the entire fibril axis longitudinally. This allows water and possibly ions to permeate through the protofilament (**Figure 5C**). P2 has the highest water permeability with an average number of water molecules of 9±2, followed by P1 (7±2) and P3 (4±3). The volumes of the newly identified cryptic cavities were 479±156 Å^3^, 314±106 Å^3^, and 304±130 Å^3^ for P1, P2 and P3, respectively. The high standard deviations are likely due to the generally high structural fluctuations observed in these sequences, in particular the abundance of conformationally flexible glycine residues.

Pocket 2 (P2) was closest to the dimerization interface of the PHF dimerization region (**Figure 5A** and **5B**). Furthermore, this pocket showed the highest druggability as predicted by fpocket (Schmidtke and Barril, 2010). P2 thus seems particularly attractive for an interaction with PHOX15. Geometrically, the radius of pocket 2 monitored along the MD simulation was 2.1 ± 0.6 Å, which compares well with that of PHOX15 of 2.4 ± 0.4 Å. This finding, together with the linearity of both PHOX15 and pocket 2, suggest possible steric complementarity. Conversely, the PHOX compounds with bulkier benzylbenzene moieties would not be sterically able to enter this channel-like pocket.

We also repeated the same analyses on a 620 ns MD simulation of the dimeric PHF structure (**Figure 5A**), obtaining similar results. Again, we identified P1, P2, and P3 on one protofilament (protofilament A) and the corresponding pockets P1*, P2*, and P3* on protofilament B. An additional pocket, P4*, was detected in protofilament B, while it was not present in protofilament A. This additional pocket may be due to the observed higher structural flexibility of protofilament B compared to protofilament A.

The radius of P2/P2* on the PHF dimer was similar to that of the single protofilament, with P2 and P2* sharing the same cylinder-like and smooth cavity with an average radius of 2.2±0.4 Å and 2.7±0.7 Å, respectively. Again, the slight difference between P2 and P2* is likely due to the higher flexibility of protomer B.

### The simulations indicate that PHOX15 disrupts the propensity of *glycine triads* to form a PHF-like assembly

Pocket 2 may have the structural features required for the binding of a small molecule as PHOX15. As described above, P2 is located at the PHF dimerization interface (**Figures 5A, 5B**). This pocket contains a cluster of three glycine residues belonging to the R3 segment (**Figure 5A**), which we have termed the *glycine triad*. According to our MD simulations of dimeric tau, these glycines and the nearby residues are likely to play a key role in assembling the PHF interface, mainly through hydrogen bonding and salt bridges. In the identified protofilament A - protofilament B contacts, a double H-bonding interaction was established through the backbone carbonyl and amide nitrogens of the central glycine residues. In addition, salt bridges between the charged side chains of Lys331 and Glu338 residues (5O3L numbering) and hydrogen bonding between the side chain of Gln336 of both protofilaments with the backbone carbonyl residue of Lys331 or Pro332 residues of the other protofilament were observed. These interactions were observed in each tau filament. Protofilament A – protofilament B contacts mediated by the *glycine triads* have occupancies in the order of 50-85%, while those mediated by the second group of residues had occupancies of 10-67% during the simulation, which highlights the key role of the central glycine residues of the two protofilaments in assembling the PHF structure. The upstream (Gly333) and downstream (Gly335) glycine residues (**Figure 5A**) form a network of H-bonds with the backbones of Pro332 and Gln336, respectively, placed on the top strand of the protofilament. These interactions have occupancies of 80% and appear to be important to stabilize the conformation of Gly334 in a PHF-like arrangement, as highlighted by the dramatic decrease in glycine-proline and glycine-glutamine intra-protofilament H-bonds in the simulation of protofilament A.

We assessed and compared the Ramachandran plots (F and ? angles) of all Gly-Gly-Gly residues along the MD simulation of dimeric PHF and single protofilament tau, both with and without bound PHOX15. As expected, the *glycine triads* in the simulation of protofilament A revealed a high degree of conformational freedom. Conversely, the conformational freedom of the *glycine triads* in the PHF dimer was very narrow, exploring only one or two states. These results are consistent with the tight H-bonding network observed in the simulation of the PHF dimer. Accordingly, we compared the F and ? angles and the hydrogen bonding patterns of the *glycine triads* typical of PHF folding in both MD simulations of the PHF dimer, protofilament A alone and protofilament A with bound PHOX15 (details about the F and ? angles and hydrogen bonding calculations are reported in the Supporting Information). As expected, the percentages of occurrence of glycine residues in the PHF-like conformation were very high (around 100%) in the MD simulation of the PHF dimer (**Table 1**). In contrast, these percentages decreased to 50-60% in the MD simulation of protofilament A alone, indicating that about half of the conformational space of the single protofilament is not prone to dimerization.

**Table 1.** Percentages of occurrence of the *glycine triads* of each tau filament adopting a PHF-like conformation. The results are reported for the MD simulations of the PHF dimer, protofilament A, and protofilament A with bound PHOX15. Numbers in the table refer to the percentages of occurrence of conformations in which the three glycine residues have F and ? angles typical of the PHF fold.

| *Tau monomers* | *PHF (%)* * | *Prot. A (%)* * | *Prot. A + PHOX15 (%)* * |
| --- | --- | --- | --- |
| *Filament 1* | 86.9 (49.2) | 12.4 (10.1) | 1.2 (0.6) |
| *Filament 2* | 97.0 (55.2) | 57.5 (38.4) | 29.6 (12.4) |
| *Filament 3* | 99.1 (55.5) | 63.5 (44.5) | 23.7 (9.6) |
| *Filament 4* | 98.1 (55.8) | 53.3 (37.4) | 37.4 (17.7) |
| *Filament 5* | 96.8 (55.4) | 11.7 (5.86) | 0.6 (0.01) |

Notes:
\* The percentages are calculated according to the formula: (number of MD frames with $\Phi$ and $\psi$ angles of the three glycine residues in the PHF conformation / total number of frames of the MD simulation) × 100. Numbers in brackets report the percentage of the glycine triads of each tau filament adopting a PHF-like conformation as well as performing the H-bond interactions of the PHF fold as described in **Figure S18**.

To assess the effect of PHOX15, we then performed docking calculations using the induced-fit docking (IFD) protocol of the Schrödinger suite 2021-1 (Sherman et al., 2006). The results of the docking of PHOX15 in pocket 2 showed that the molecule can accommodate itself longitudinally in P2 resulting in favorable hydrogen bonding and hydrophobic interactions. Starting from the docking complex, an 800 ns MD simulation of the protofilament A in complex with PHOX15 showed a further decrease in the percentages of occurrence of PHF-like conformations, which were significantly lower than those observed in the simulations of the PHF dimer and of the protofilament A alone (see the percentages of occurrence of tau conformations with F and ? angles typical of the PHF-like conformation reported in **Table 1**. Similar conclusions could be drawn by including in the analysis the % of occurrence of tau conformations with the hydrogen bond network present in the PHF fold, which decreased significantly in the simulation of the protofilament A – PHOX15 complex (**Table 1**). Altogether, the results of the simulations suggest that PHOX15 could bind pocket 2 of protofilament A and significantly reduces the ability of this protofilament to adopt a PHF-like conformation, mainly by affecting the conformation of the *glycine triads*. Docking and MD simulations of PHOX15 in the PHF dimer showed that PHOX15 is no longer able to influence its conformation after the formation of the dimer (data not shown). This finding is consistent with PHOX15 exerting an effect at early stages of tau oligomerization.

## DISCUSSION

Tau aggregation and impaired microtubule interaction are believed to play a central role in the development of AD and other tauopathies. Therefore, drugs that inhibit tau aggregation or increase the interaction of tau with microtubules could be a promising approach to combat tau-induced neurodegeneration. In fact, several drug candidates aiming to inhibit tau aggregation, reduce tau phosphorylation, or decrease tau expression are under investigation (Soliman et al., 2022). However, mechanism-based drug assays are difficult to establish due to the lack of cell-based models that allow to monitor tau aggregation and effects on tau-microtubule interaction of the full-length tau protein. While previously “biosensor cells” were generated to probe pathological tau seeds, these cells are based on the expression of only a fragment of the tau repeat domain that does not contain additional regulatory regions such as the proline-rich region (Hitt et al., 2021; Holmes et al., 2014). The presence of additional regions is of particular importance for the analysis of the role of tau in the disease process, because the tau region involved in mediating tau aggregation and its binding to microtubules overlaps, and regions flanking the microtubule-binding repeats of tau can modulate both activities (Fitzpatrick et al., 2017; Kadavath et al., 2015; Niewidok et al., 2016). Therefore, approaches aiming to inhibit tau aggregation run the risk of also negatively affecting the physiological interaction of tau with microtubules.

In addition, increasing evidence suggests that earlier stages of tau aggregation, particularly soluble oligomeric tau species that precede the formation of neurofibrillary tangles (NFTs), are the toxic species (Cowan et al., 2010; Spires et al., 2006; Wittmann et al., 2001). Indeed, NFTs may actually be neuroprotective by sequestering tau (Cowan and Mudher, 2013), which would require the development of tau aggregation inhibitors that selectively inhibit the early stage of tau aggregation without affecting the existing NFTs. Experimental evidence also suggests that disease-like hyperphosphorylation of tau and microtubule binding play a crucial role in the regulation of tau toxicity (Chatterjee et al., 2009; Fath et al., 2002), implying that it is important to differentiate the effect of potential drugs on tau aggregation, tau phosphorylation and tau-microtubule interaction in order to develop approaches that positively modulate certain features of tau toxicity.

The multiple facets of tau’s involvement in the neurodegenerative cascade also make tauopathies an attractive target for a polypharmacological approach, particularly using computational methods that offer the ability to predict the activity profile of ligands in the iterative design and optimization steps of a preclinical drug discovery project (Rastelli and Pinzi, 2015). Along these lines, it would be quite attractive to have a drug that simultaneously inhibits early tau aggregate formation and reduces tau phosphorylation at disease-relevant sites, thereby restoring physiological tau-microtubule interaction and regulation of axonal microtubule polymerization.

In this work, we: (1) developed a quantitative live-cell imaging approach to analyze tau-microtubule interaction in axon-like processes of a tauopathy model; (2) we used the approach to screen a set of candidate molecules that, according to chemoinformatic analysis, inhibit tau aggregation and disease-associated kinases; (3) we identified one compound, PHOX15, that inhibited an early state of tau aggregation *in vitro*, restored tau-microtubule interaction of an aggregation-prone tau construct in neurons, and increased tau-microtubule interaction by reducing tau phosphorylation at disease-relevant sites; and (4) we found cryptic pockets in tau protofilaments with channel-like features through molecular dynamic simulations where PHOX15 might bind and decrease the ability to adopt a PHF-like conformation.

Our data indicate that PHOX15 normalizes the tau-microtubule interaction through two distinct mechanisms (**Figure 5D**). On the one hand, it increases the interaction of an aggregation-prone tau construct with microtubules to a physiological level, most likely by reducing the amount of soluble tau oligomers. This activity might be mediated by PHOX15 binding to cryptic channel-like pockets crossing tau protofilaments that we have identified through MD simulations, thereby disrupting tau oligomerization. In this regard, this type of interaction is favorable as it would interfere with tau aggregation but would not compete with tau binding to microtubules. On the other hand, PHOX15 reduces tau phosphorylation at disease-relevant sites, particularly in the proline-rich region, which contains a cluster of phosphorylation sites that can be phosphorylated by GSK3*β*. Indeed, most sites in this region that show more than two-fold reduced phosphorylation are predicted phosphorylation sites for GSK3*β* (i.e., Ser235, Ser237, Thr231, Ser195, Thr181, Thr217, Ser198, Ser199, Ser210; NetPhos-3.1b prediction). Analysis of kinome activity showed that PHOX15 changes the activity of several kinases in addition to GSK3*β*, and it has yet to be shown that this does not have any undesirable side effects that would require further drug optimization.

Our PC12 cell data show that the aggregation-prone tau construct does not form insoluble tau aggregates in the cells and cells transiently expressing the TauΔK280 construct were negative for staining with Amytracker 680, a probe that binds to amyloid fibrils with at least eight repetitive beta-sheets (Ebba Biotech, Solna, Sweden) (data not shown). Lentiviral infection of DRG neurons also allowed the effect of prolonged presence of aggregation-prone tau to be determined, and we observed Amytracker-positive tau aggregates after three weeks of continuous expression. This indicates that already the formation of small soluble tau oligomers has a significant impact on the interaction of tau with microtubules, and we observed an approximately 10% reduced tau-microtubule interaction. The data also show that PHOX15 binds to small tau oligomers and effectively increases the availability of tau to interact with microtubules.

Although PHOX15 needs further testing in a systemic setting, many previous drug studies were initiated without a thorough mechanism-based analysis of their functions, which is particularly critical for a complex target like tau with many interaction partners and regulation by various post-translational modifications (Brandt et al., 2020; Morris et al., 2011). We are confident that our novel cell model to study the effect of drugs on the modulation of tau aggregation and tau-microtubule interaction and the identification of the small molecule PHOX15 as a promising polypharmacological drug candidate will help advance approaches that specifically target the key traits of tauopathies. Furthermore, we are confident that the simulations provide the basis for a structure-based optimization of the activity of these compounds as well as the rational design of new anti-aggregation tau ligands.

## METHODS

### Expression vectors and virus preparations

Prokaryotic expression plasmids were based on human adult tau (Tau441wt) in a pET-3d vector (Brandt and Lee, 1993). Eukaryotic expression plasmids for tau variants were based on human adult tau (Tau441wt) or fetal tau (Tau352wt) with an amino-terminally fused PAGFP tag in a pRc/cytomegalovirus (CMV) expression vector (Gauthier-Kemper et al., 2011). The ΔK280 deletion was introduced into the expression plasmids by site-directed mutagenesis using phosphorylated primers (5’AAGCTGGATCTTAGCAACGTC and 5’ATTAATTATCTGCACCTTCCCGCC) with Platinum SuperFi DNA Polymerase (ThermoFisher Scientific, USA). The phosphoblocking tau mutant (Tau352Ala) was constructed by changing the codons for Ser198, Ser199, Ser202, Thr231, Ser235, Ser396, Ser404, Ser409, Ser413, and Ser422 in GCT (alanine) as previously described (Eidenmuller et al., 2000). Lentiviral vectors for Tau441wt and TauΔK280 with amino-terminally fused PAGFP tag were constructed in L22FCK(1.3)GW (provided by P. Osten, Northwestern University, Chicago, IL) containing the neuron-specific promoter α-CaMKII (Dittgen et al., 2004). Sequences introduced by PCR were verified by DNA sequencing (Seqlab-Microsynth, Göttingen, Germany). For the production of lentivirus, 293FT human embryonic kidney cells (Thermo Fisher Scientific, USA) were transfected with the expression vector and two helper plasmids and viral particles from the supernatant were concentrated by ultracentrifugation as previously described (Bakota et al., 2012).

### Cell lines

PC12 cells were cultured in serum-DMEM (DMEM supplemented with 10% fetal bovine serum and antibiotics (100 U/ml penicillin and 100 µg/ml streptomycin)), at 37°C with 10% CO_2_ in a humidified incubator and transfected with Lipofectamine 2000 (Thermo-Fisher Scientific, USA) as previously described (Fath et al., 2002). After transfection, the medium was replaced with serum-reduced DMEM (DMEM supplemented with 1% fetal bovine serum and antibiotics) and the cells were neuronally differentiated for 4 days by the addition of 100 ng/ml 7S mouse NGF (Alomone Labs, Israel).

### Primary neuronal cultures

C57BL/6J mice were kept and killed in accordance with the German animal care regulations based on the FELASA guidelines. Three-month- or one-year-old mice were sacrificed by cervical dislocation. The spine was quickly dissected and the spinal cord exposed. Dorsal root ganglia (DRGs) were extracted, dissociated and enzymatically digested essentially as previously described (Sleigh et al., 2016). The resulting suspension, containing isolated DRG neurons, was plated in DRG culture medium (Neurobasal A supplemented with 2% B-27, 1% fetal bovine serum, 1% horse serum, 20 µM L-glutamine, 0.1% *β*ME, 100 µg/ml Primocin®) supplemented with 100 ng/mL 7S mouse NGF. Infectious lentiviral particles were applied on day 3. After 6 hours of incubation, the medium was replaced with fresh medium containing NGF, and incubation was continued at 37°C with 5% CO_2_ in a humidified incubator. Live cell imaging was performed on day 10 *in vitro*.

### Chemicals and cell treatments

The 2-phenyloxazole (PHOX) derivatives (Di Paolo et al., 2019) were prepared as 20 mM stock solutions in DMSO and were stored at -20°C in light-protected Eppendorf cups.

### Measurements of neuronal viability

Metabolic activity and viability were assessed using a combined MTT/LDH assay. PC12 cells were cultured in 96-well plates at 10,000 cells per well in 50 μl serum-reduced DMEM supplemented with 100 ng/ml 7S mouse NGF. After 48 hours, test compounds at final concentrations of 0, 6.25, 12.5, 25, 50 and 100□µM were added in a further volume of 50 µl and incubation was continued for 20 hours. For LDH measurements, 50µl from each well was transferred to a new 96-well plate and 50µl LDH-reagent (4 mM iodonitrotetrazolium chloride (INT), 6.4 mM beta-nicotinamide adenine dinucleotide sodium salt (NAD), 320 mM lithium lactate, 150 mM 1-methoxyphenazine methosulfate (MPMS) in 0.2 M Tris-HCl buffer, pH 8.2) was added. The plate was shaken for 10 seconds and incubated in the dark for 10 minutes. Absorbance was measured at 490 nm using a ThermoMax Microplate Reader operated with SoftMaxPro Version 1.1 (Molecular Devices Corp., Sunnyvale, USA.). Measurements were normalized to a positive control to which 1% Triton X-100 had been previously added. For MTT measurements, 50 µl of MTT-reagent (2 mg/ml MTT (3,(4,5.dimethylthiazol-2-yl)2,5-diphenyltetrazolium bromide in prewarmed serum-reduced DMEM) was added to each of the remaining wells. Cells were incubated for 2 hours at 37°C and the reaction was stopped by the addition of 50 μl lysis buffer (20% (wt/vol) sodium dodecyl sulfate in 1:1 (vol/vol) N,N-dimethylformamide/water, pH 4.7). After overnight incubation at 37°C, optical densities at 570 nm were determined. Measurements for MTT conversion were normalized to optical densities of the negative control wells. All experiments were performed in triplicates in two independent plates.

### Purification of recombinant tau protein

pET-3d-tau plasmids were transformed into Escherichia coli BL21(DE3)pLysS cells for expression, and the cells were grown, induced and harvested as previously described (Studier et al., 1990). Tau was purified from the cell extract of 200 ml of culture by sequential anion exchange and phosphocellulose chromatography as previously described (Brandt and Lee, 1993). Tau protein eluate was dialyzed against PBS containing 2 mM MgCl_2_, concentrated with Vivaspin® (15R, 2,000 MWCO, Sartorius, UK) and adjusted to 1 mM DTT. Protein concentrations were determined by densitometry of Coomassie Brilliant Blue-stained gels using bovine serum albumin as a standard.

### Tau aggregation assay

Assays were performed with 0.5 mg/ml recombinant tau protein in 30 mM MOPS/NaOH (pH 7.4) containing 200 µg/ml heparin and 1 mM 4-(2-aminoethyl)benzenesulfonyl fluoride (AEBSF, Applichem, Darmstadt, Germany) in an assay volume of 25 µl essentially as previously described (Eidenmuller et al., 2000). Incubation was at 37 °C for the indicated times and concentrations. In order to quantify the amount of aggregates, the mixture after the assembly reaction was centrifuged for 1 h at 100,000×g and 4 °C. One third of the supernatant and pellet fractions were separated by SDS-PAGE with 15% acrylamide, stained with Coomassie Brilliant Blue and quantitated by densitometry.

### Electron microscopy

For analysis by transmission electron microscopy (EM), aggregation assays were performed with 0.4 mg/ml recombinant TauΔK280 in an assay volume of 25 µl for 24 hrs at 37°C. To test for drug induced disassembly, PHOX15 or carrier were added in 10% of the volume, mixed, and incubation continued for another 24 hrs. The samples were sonicated for 2 min twice with a pause of 1 min. Pioloform-coated and glow-discharged grids were floated on a drop of the sample for 10 min, washed with water, negatively stained with 1% uranyl acetate, and dried. EM was performed with a Zeiss 10CR electron microscope at 60 kV.

### Immunoblotting

After separation by SDS–PAGE, proteins were transferred to Immobilon-P PVDF membranes (Millipore, USA), followed by immunoblotting. Detection employed PHF1 antibody and peroxidase-conjugated donkey anti-mouse secondary antibody (Jackson ImmunoResearch Laboratories, Inc., USA). Protein bands were detected using enhanced chemiluminescence with SuperSignal West Dura extended duration substrate (Thermo Fisher Scientific, USA) according to the manufacturer’s protocol. Quantification of the blots was carried out with Gel-Pro Analyzer 4.0 (Media Cybernetics L.P., USA) or with FusionCapt Advance (Vilber Lourmat, France).

### Live-cell imaging and Fluorescence Decay After Photoactivation (FDAP)

For FDAP experiments, wild-type tau or TauΔK280 expressing cells were plated on 35-mm poly-L-lysine and collagen-coated (for PC12 cells) or laminin-coated (for DRG neurons) glass-bottom culture dishes (MatTek, USA). PC12 cells were neuronally differentiated after transfection by medium exchange for serum-reduced DMEM containing 100 ng/mL 7S mouse NGF. Cultivation was continued for 4 days with medium exchange for serum-reduced DMEM containing NGF and without phenol red one day prior to live imaging. Treatment of the cells was performed 20 hours prior to live imaging by addition of the respective compound (or DMSO for carrier control) in the desired concentration. Live cell imaging for photoactivation experiments was essentially performed as described previously (Conze et al., 2022) using a laser scanning microscope (Nikon Eclipse Ti2-E (Nikon, Japan)) equipped with a LU-N4 laser unit with 488-nm and 405-nm lasers and a Fluor 60× ultraviolet-corrected objective lens (NA 1.4) enclosed in an incubation chamber maintaining 37°C and 5% CO_2_. Photoactivation of a 6 μm long neurite segment was performed with a 405-nm laser. A set of consecutive image series (time stack) was obtained at a frequency of 1 frame/s, and 112 frames were collected per activated cell at a resolution of 256×256 pixels. Effective diffusion constants and fraction of microtubule-bound tau were determined by fitting the fluorescence decay data from photoactivation experiments using a one-dimensional diffusion model function for FDAP, as previously described (Weissmann et al., 2009). For staining with optotracer, Amytracker 680 was diluted 1:500 in the culture medium. After 30 minutes of incubation, cells were imaged with a 453-nm laser.

### Proteome and phosphoproteome analysis

PC12 cells were neuronally differentiated by culture for 4 days in serum-reduced DMEM containing 100 ng/ml 7S mouse NGF. 20 hours prior to sample preparation, cells were treated with 25 µM PHOX15 or carrier (0.125% DMSO). Cells were washed in ice-cold PBS, incubated in lysis buffer (8M urea in 50mM Tris/HCl, pH 7.8) supplemented with Phos-Stop tablets (Roche Diagnostics GmbH, Germany) and sonicated for 5 cycles of 5 s each at 10% amplitude (Branson Digital Microtip Sonifier 250-D, Connecticut, USA). Samples were cleared by centrifugation at 4°C and 23,000×g for 30 min and protein concentration determined using Pierce(tm) BCA Protein Assay (Thermo Fisher Scientific, USA). 0.2 μg/μl α-casein were added to a protein amount of 1.2 mg, and the reduction and alkylation were carried out in lysis buffer containing 15 mM iodoacetamide and 5 mM DL-dithiothreitol. Proteins were digested for 18 h with trypsin/Lys-C Mix (Promega Corporation, USA) and 10 μg of each sample was used for proteome analysis. For phosphoenrichment, remaining samples were desalted using Sep-Pak® Classic C18 cartridges (Waters, Ireland) that had been prewashed and equilibrated with 5 ml methanol, 5 ml 80% acetonitrile and 2×5 ml 0.5 % formic acid. After sample application, the Sep-Pak was washed once with 5 ml of 0.5 % formic acid and eluted with 4 ml of 80% acetonitrile/0.5 % formic acid. The eluate was lyophilized and enriched with a High-Select(tm) TiO2 Phosphopeptide Enrichment Kit (Thermo Fisher Scientific, USA). For proteome and phosphoproteome analysis, samples were taken from a PepMap C18 easy spray column (Thermo Fisher Scientific, USA) with a linear gradient of acetonitrile from 10–35% in H_2_O_2_ with 0.1% formic acid for 180 min at a constant flow rate of 25 nl/min.. MS analysis was performed as previously described (Schoppe et al., 2020). The *.raw data files were analyzed with PEAKS Online software (Bioinformatic Solutions Inc, Canada). PEAKS Q (de-novo-assisted quantification) analysis was used for data refinement with mass correction, de-novo sequencing and de-novo-assisted database search, and subsequent label-free quantification. The search engine was applied to *Rattus norvegicus* *.fasta databases. MS/MS searches were performed using a mass tolerance of 10 ppm parent ions and a mass tolerance of 0.2 Da fragments. Trypsin with up to two missing cleavage points was selected as the cleavage enzyme. Carbamidomethylation modification was chosen as the fixed modification and the oxidation of methionine, acetylation of lysine and phosphorylation of serine, threonine and tyrosine were chosen as the variable modifications. A maximum of three variable modifications were allowed per peptide. Normalization to the total ion current level for each sample was performed. ANOVA test was used to calculate the significance level for each protein, outliers were removed, and the top three peptides were used for protein signal quantification where it was possible. The peptide identification was considered valid at a false detection rate of 1% (q-value < 0.001) (maximum delta Cn of the percolator was 0.05). The minimum length of acceptable identified peptides was set to six amino acids. Each condition was analyzed in triplicate. All proteins were assigned their gene symbol via the Uniprot knowledge database (http://www.uniprot.org/). Tau interaction partners selected for representation on a volcano plot were based on previously described region-specific interactors (Brandt et al., 2020). Normalized phosphoproteomics data revealed phosphopeptides that were down-regulated after treatment with PHOX15, gene symbols associated with these phosphopeptides were used for kinase enrichment analysis via KEA3 application (Kuleshov et al., 2021). Mean rank and the number of substrates that were associated with fifteen highest-ranking kinases were used for representation.

### Statistical analysis

Statistical analysis was carried out with GraphPad Prism v8.0.1 (GraphPad Software, USA). All data sets were tested for normality using D’Agostino-Pearson and Shapiro-Wilk test. If necessary, data sets were log transformed in order to enable further statistic testing. Statistical outliers were identified using the ROUT method. The homogeneity was assessed using Levene’s test. An unpaired two-tailed t-test was used to compare two datasets. In the case of unequal variances, the Welch’s correction was applied. A one-way ANOVA was performed to compare more than two data sets followed by Dunnett’s post-hoc test. All statistical values are expressed as mean ± SEM.

## ACKNOWLEDGMENT

We are grateful to Hilmar Bading, Department of Neurobiology and Interdisciplinary Center for Neurosciences, University of Heidelberg, for providing the opportunity to carry out the electron microscopy work in his laboratory. This work was supported by the Deutsche Forschungsgemeinschaft (DFG BR1192/14-1 to RB), Fondo di Ateneo per la Ricerca (FAR 2019, DR496/2019 to GR) and European Union’s Horizon 2020 research and innovation program H2020-MSCAITN-2019-EJD - grant agreement no: 860070 (NB, AS, DP and RB).

## AUTHOR CONTRIBUTIONS

Luca Pinzi: Investigation, Writing - Original Draft; Christian Conze: Investigation, Writing - Original Draft; Nicolo Bisi: Investigation, Writing - Review & Editing; Gabriele Dalla Torre: Investigation, Writing - Review & Editing; Nanci Abreu: Investigation, Writing - Review & Editing; Nataliya I. Trushina: Investigation, Writing - Review & Editing; Ahmed Soliman: Investigation, Writing - Review & Editing; Andrea Krusenbaum: Investigation, Writing - Review & Editing; Maryam Khodaei Dolouei: Investigation, Writing - Review & Editing; Andrea Hellwig: Investigation, Writing - Review & Editing; Michael S. Christodoulou: Resources; Daniele Passarella: Resources; Lidia Bakota: Writing - Original Draft, Supervision; Giulio Rastelli: Conceptualization, Writing - Original Draft, Supervision, Funding acquisition; Roland Brandt: Conceptualization, Writing - Original Draft, Supervision, Funding acquisition.

## Notes

### Competing Interest Statement

The authors have declared no competing interest.

